# Palaeoproteomic investigation of an ancient human skeleton with abnormal deposition of dental calculus

**DOI:** 10.1101/2023.08.15.553159

**Authors:** Yoko Uchida-Fukuhara, Shigeru Shimamura, Rikai Sawafuji, Takumi Nishiuchi, Minoru Yoneda, Hajime Ishida, Hirofumi Matsumura, Takumi Tsutaya

**Affiliations:** Department of Oral Morphology, Faculty of Medicine, Dentistry and Pharmaceutical Sciences, Okayama University, Okayama 700-8525, Japan; Research Center for Integrative Evolutionary Science, The Graduate University for Advanced Studies (SOKENDAI), Kanagawa 240-0193, Japan; Institute for Extra-Cutting-Edge Science and Technology Avant-Garde Research (X-STAR), Japan Agency for Marine-Earth Science and Technology (JAMSTEC), Yokosuka 237-0061, Japan; Department of Human Biology and Anatomy, Graduate School of Medicine, University of the Ryukyus, Okinawa 903-0215, Japan; Research Center for Experimental Modeling of Human Disease, Kanazawa University, Nanazawa 920-8640, Japan; The University Museum, The University of Tokyo, Tokyo 113-0033 Japan; Mt. Olive Hospital, Okinawa 903-0215, Japan; School of Health Sciences, Sapporo Medical University, Hokkaido 060-8556, Japan; Biogeochemistry Research Center (BGC), Japan Agency for Marine-Earth Science and Technology (JAMSTEC), Yokosuka 237-0061, Japan

## Abstract

Detailed investigation of extremely severe pathological conditions in ancient human skeletons is important as it could shed light on the breadth of potential interactions between humans and disease etiologies in the past. Here, we applied palaeoproteomics to investigate an ancient human skeletal individual with severe oral pathology, focusing our research on bacterial pathogenic factors and host defense response. This female skeleton, from the Okhotsk period (i.e., 5th–13th century) of Northern Japan, poses relevant amounts of abnormal dental calculus deposition and exhibits oral dysfunction due to severe periodontal disease. A shotgun mass-spectrometry analysis identified 81 human proteins and 15 bacterial proteins from the calculus of the subject. We identified two pathogenic or bioinvasive proteins originating from two of the three “red complex” bacteria, the core species associated with severe periodontal disease in modern humans, as well as two additional bioinvasive proteins of periodontal-associated bacteria. Moreover, we discovered defense response system-associated human proteins, although their proportion was mostly similar to those reported in ancient and modern human individuals with lower calculus deposition. These results suggest that the bacterial etiology was similar and the host defense response was not necessarily more intense in ancient individuals with significant amounts of abnormal dental calculus deposition.

## Introduction

Ancient human skeletons sometimes show abnormal and extremely severe pathological conditions that could be rarely observed in modern human populations^1,2^. Such extreme cases could be considered “natural experiments” that highlight both human resilience and vulnerability to disease in the absence of modern medical interventions^3,4^. Humans and pathogens coevolved and various ancient pathogens are not equivalent to their contemporary descendants^5,6^. Ancient severe pathological conditions that cannot be seen today could have existed due to the lack of modern medical interventions or different bacterial etiologies. Detailed investigation of these extreme cases would be important as they shed light on the breadth of potential interactions between humans and diseases, and reveal differences between past disease etiologies and present-day pathogens.

In this study, we used palaeoproteomics to investigate the etiology of and host resilience to periodontal disease in an ancient human skeleton showing abnormal deposition of dental calculus with severe periodontal disease. Dental calculus is a calcified oral plaque that promotes periodontal disease^7^ and is habitually removed in modern dental care. In contrast, abnormal depositions of dental calculus, where a large calculus deposition entirely covers the occlusal surface of at least one tooth, could be occasionally observed in ancient human skeletons. Such examples include a late Saxon skeleton from Nottinghamshire, UK^8^, and the subject of this study, an Okhotsk skeleton from Hokkaido, Japan^9^. Dental calculus entraps and preserves microparticles, DNA, and proteins originating from the environment, host, microbiome, and diet. Therefore, dental calculus provides molecular clues to help understand the lifeways of the host, pathological conditions, and disease etiology in the past^10,11^. Analyzing abnormally deposited dental calculus can further reveal the pathogenic cause of oral pathology and the defense response of the host.

Palaeoproteomics of dental calculus, applied in this study, is an effective method for investigating both the etiology of and host responses to ancient periodontal disease^12–14^. Proteins are functional agent, and their expression differs in response to pathological conditions. These pieces of evidence, revealing information on functional oral pathologic processes, could not be obtained solely by DNA analysis, which could only reveal the presence of certain taxa in analyzed specimens. The paleoproteomic analytic potential of dental calculus for studying health and diseases in the past has not been fully exploited (however, see references^12–14,15^) despite successful applications in studies aiming at dietary reconstruction^16–21^.

By applying palaeoproteomics to abnormally deposited dental calculus from a skeletal individual with severe periodontal disease, we aimed at answering i) whether the pathogenic factors associated with the severe periodontal disease in this individual differed from modern and ancient human individuals with lower calculus deposition, and ii) to what extent the extreme oral pathological conditions caused pathological stress to the host.

### The subject individual, HM2-HA-3

HM2-HA-3 is a female skeleton, aged 34–54 years at death, excavated in 1992 from the Hamanaka 2 site (Figure 1) on Rebun Island, Hokkaido, Japan^22^. The most notable feature of this individual is the abnormal deposition of large amounts of dental calculus (Figure 1^9^). The morphological characteristics of this individual have been previously described in detail^9^. Briefly, most skeletal elements of HM2-HA-3 were missing; only a part of the cranium, an upper limb, and trunk bones were present, though the mandible and maxilla, including erupted teeth were well-preserved. Heavy deposits of dental calculus were present, especially on the right side of the dentition. These calculus deposits are predominantly located above the cementoenamel junction, a feature of supragingival calculus. These deposits were primarily found on the right upper second and third molars (Figure 1). The occlusal surfaces of these molars are completely covered by calculus deposits and present a non-smooth surface.

**Figure 1.**
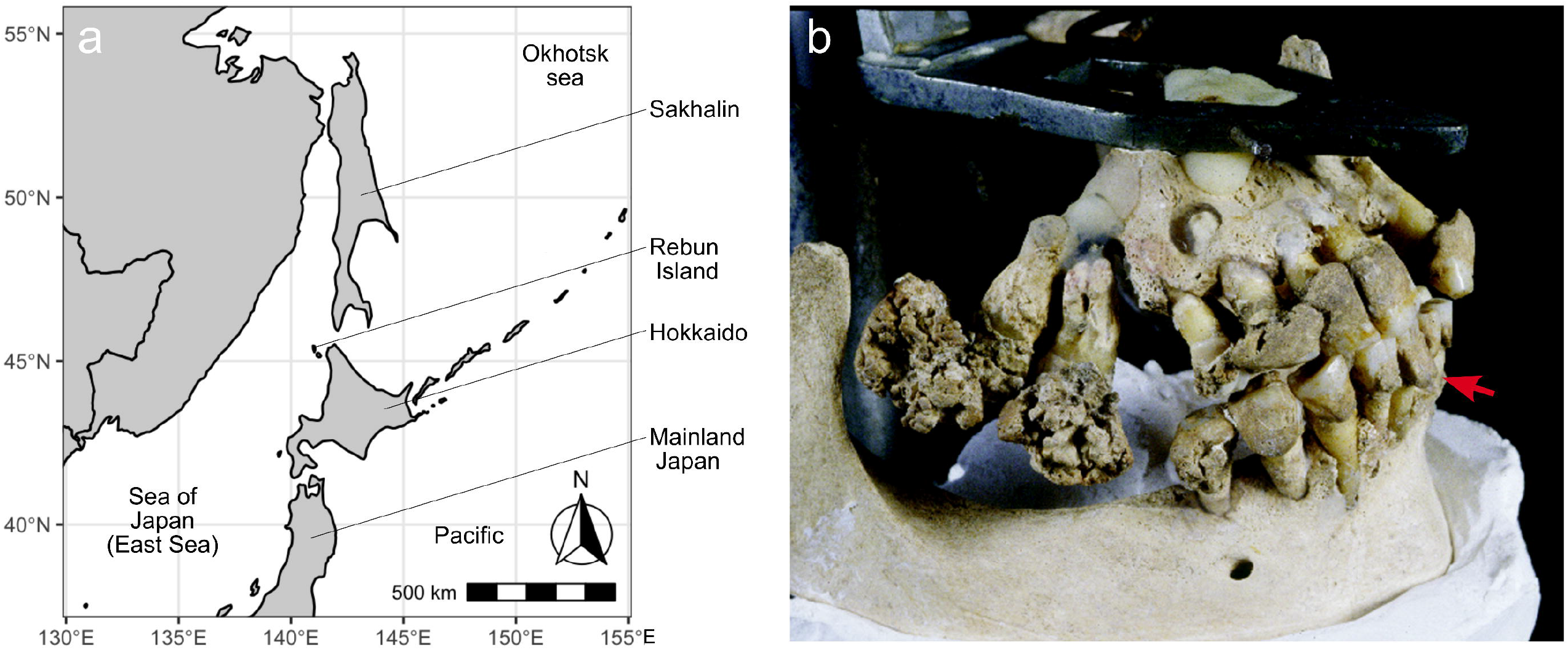
a) Map of Rebun Island and Hamanaka 2 site. b) Right buccal aspect of the HM2-HA-3 maxilla and mandible. A red arrow indicates the sampled calculus (i.e., from the lower right permanent first incisor).

HM2-HA-3 also exhibits extreme oral pathological conditions. Caries are not present in any of the remaining teeth but HM2-HA-3 presents apical lesions with cementum hyperplasia, rounded cavities in the root apex, and severe periodontal disease including resorption of the alveolar process^9^. Periodontitis-related horizontal alveolar bone resorption was prominent in HM2-HA-3, and the mandibular right molars had been completely lost with severe resorption of the crest. This individual would likely have suffered from periodontal disease since the relatively early stages of her life, when the right side of her jaws would have become almost completely unusable for masticatory function^9^. As a result, HM2-HA-3 showed severe tooth wear on her left teeth, which were not covered by calculus. Furthermore, alveolar bone resorption at the root branch was observed on the upper right side, suggesting the presence of endodontic-periodontal disease. Abnormal calculus deposition would have facilitated periodontal tissue collapse in the same region. Taken together, these conditions show that normal masticatory function would have been impaired in this individual.

HM2-HA-3 was found in an archaeological site belonging to the Okhotsk culture. The Okhotsk culture was distributed along southern Sakhalin Island, the northeastern coast of Hokkaido, and the Kuril Islands during the 5th–13th centuries^23^. The Okhotsk people predominantly subsisted on fishing, and it is estimated that marine foods comprised more than 80% of their dietary protein intake^24,25^. Although a few crop remains have been excavated from Okhotsk sites^26^, it is believed that plant horticulture was not practiced in the Okhotsk culture^23^. Because of their low carbohydrate intake, the caries rate of Okhotsk people was remarkably lower than in Jomon hunter-gatherers^27^. Physical anthropological measures of oral health, such as the frequency of linear enamel hypoplasia, in the Okhotsk people were generally better than in the Jomon hunter-gatherers of mainland Japan^28^. Even though, no other Okhotsk human skeletons show such an abnormal calculus depositions seen in HM2-HA-3^9^.

## Results

### Chronological age and diet

Elemental and isotopic results of the rib bone collagen sample from HM2-HA-3 are shown in Table 1. Bone collagen extracted from the rib of HM2-HA-3 showed acceptable %C (44.5%), %N (16.4%), and C/N ratio (3.17)^44,45^, suggesting good molecular preservation of this individual.

**Table 1.** Results of stable isotope analysis and radiocarbon measurement. Previously reported data from other skeletal individuals from the Hamanaka 2 site are also shown.

| ID | sex | Age (y) | Element | %C | %N | $\delta^{13}\text{C}$ | $\delta^{15}\text{N}$ | C/N | $^{14}\text{C}$ age (BP) | Reference |
| --- | --- | --- | --- | --- | --- | --- | --- | --- | --- | --- |
| 1480 | M | 40–50 | Skull | 43.9 | 15.1 | -13.2 | 19.0 | 3.4 | – | Naito et al., 2010<sup>24</sup> |
| 1496 | F | 30–40 | Skull | 43.8 | 15.7 | -12.9 | 18.6 | 3.3 | – | Naito et al., 2010<sup>24</sup> |
| NAT002 | F | 40–49 | – | 41.8 | 15.0 | -12.9 | 19.3 | 3.2 | – | Okamoto et al., 2016<sup>49</sup> |
| HM2-HA-3 | F | 35–54 | Rib | 44.5 | 16.4 | -13.0 | 19.3 | 3.2 | $1777 \pm 37$ | This study |

The calibrated radiocarbon age of HM2-HA-3 was 485–760 cal AD with 95.4% posterior probability and 565–678 cal AD with 68.3% posterior probability. Considering the chronology of the Hamanaka 2 site^46^, this age falls in the earlier Okhotsk period. The *δ*^13^C and *δ*^15^N values of bone collagen from HM2-HA-3, which mostly represent protein dietary components assimilated during ∼10 years before death^47,48^, were -13.0‰ and 19.3‰, respectively. These isotope ratios are shown in Figure 2 along with the previously reported values from other human skeletons excavated at the Hamanaka 2 site^24,49^ and faunal bones excavated from another Okhotsk site (Moyoro site^25^). These comparisons showed that most dietary proteins of HM2-HA-3 were obtained from marine foods and there were no apparent differences in dietary food sources between HM2-HA-3 and other Okhotsk individuals excavated from the Hamanaka 2 site (Figure 2).

**Figure 2.**
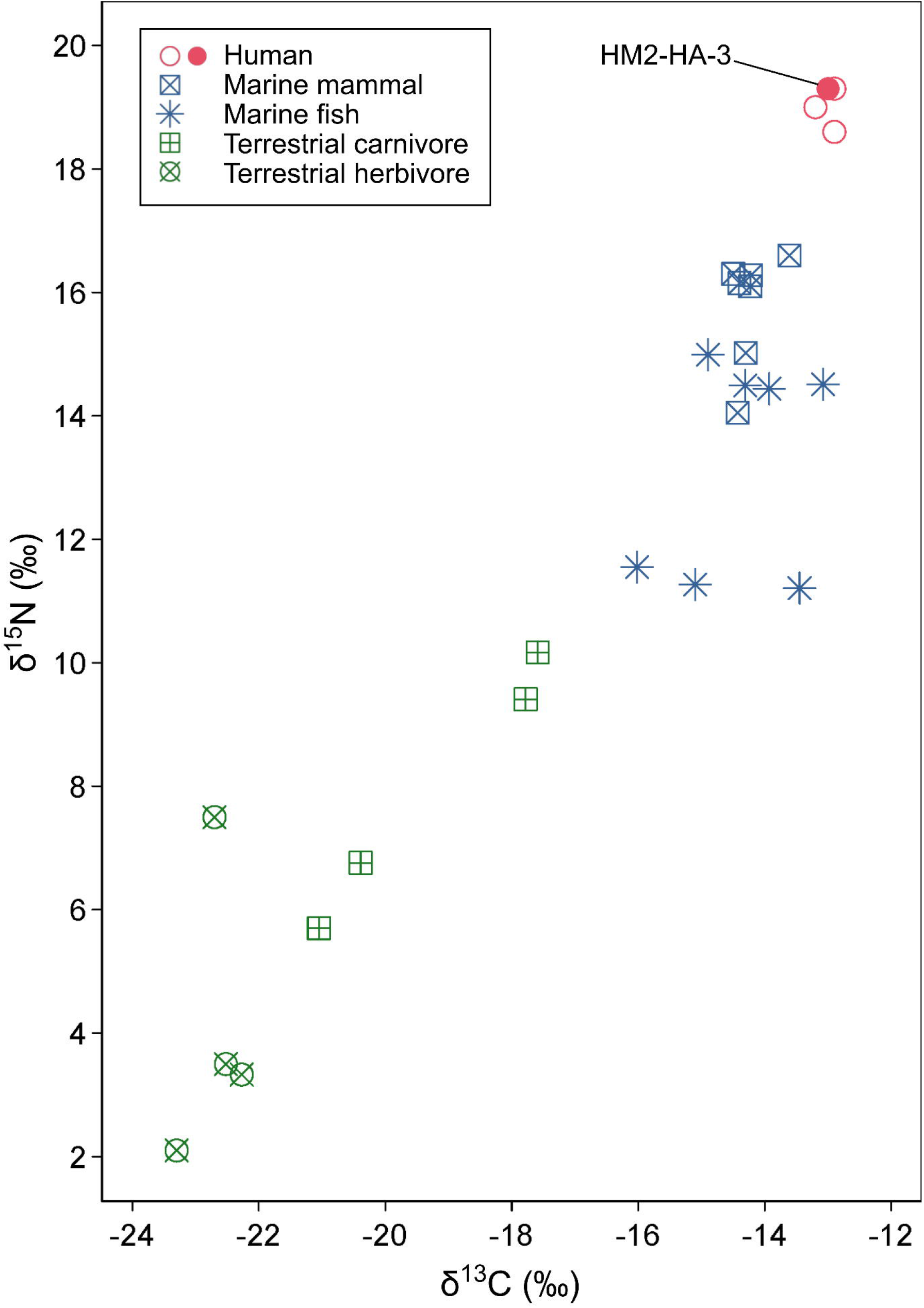
Carbon and nitrogen stable isotopic results of faunal and human bone collagen.

### Dental calculus proteome

We identified a total of 96 protein groups from the dental calculus of HM2-HA-3, excluding keratins and common laboratory contaminants. Of these, 81 and 15 protein groups originated from humans (Table 2) and bacteria (Table 3), respectively. The calculus displayed a high (i.e., 92.1%) OSSD score, suggesting good protein preservation^19^. The peptide deamidation rates, the approximate proxy for ancient protein authenticity^50,51^, derived from the four fractions ranging between 38.7%–54.8% and 30.7%–37.7% for asparagine and glutamine in human proteins, respectively (Supplementary Table S1). As the deamidation rate of modern proteins is typically below 20%, the human proteins identified in the dental calculus of HM2-HA-3 would originate from ancient times^12^.In contrast, bacterial proteins showed lower asparagine and glutamine deamidation rates (4.9%–23.2% and 4.2%–24.0%, respectively) (Supplementary Table S1). The number of asparagine and glutamine residues in the identified bacterial proteins was below 8, the precise deamidation rates could thus not be calculated.

**Table 2.**
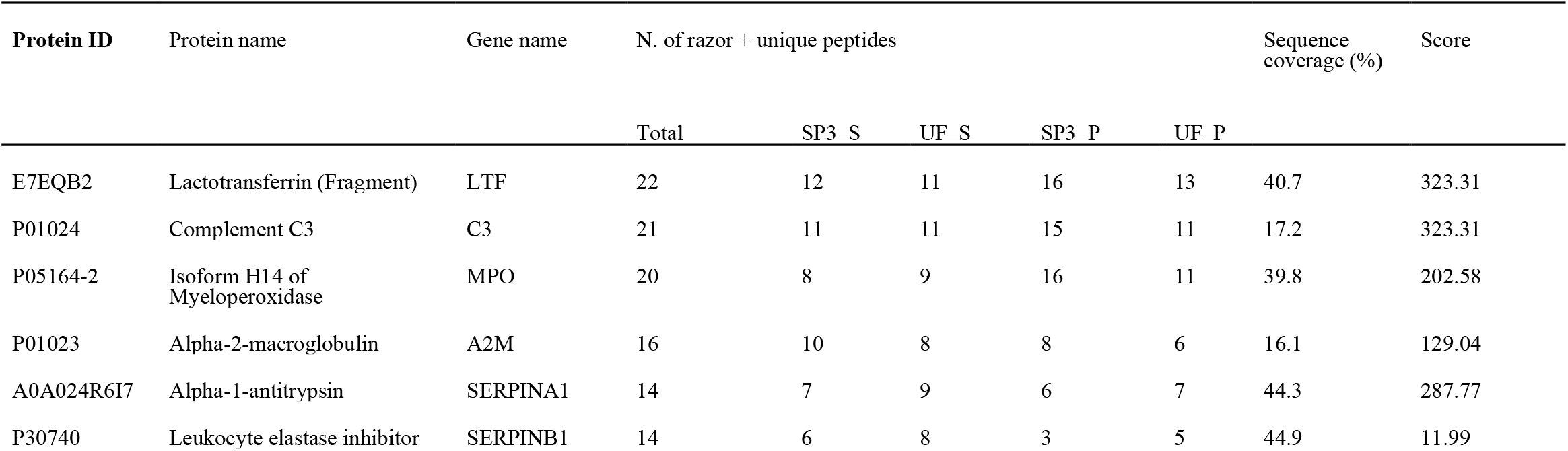
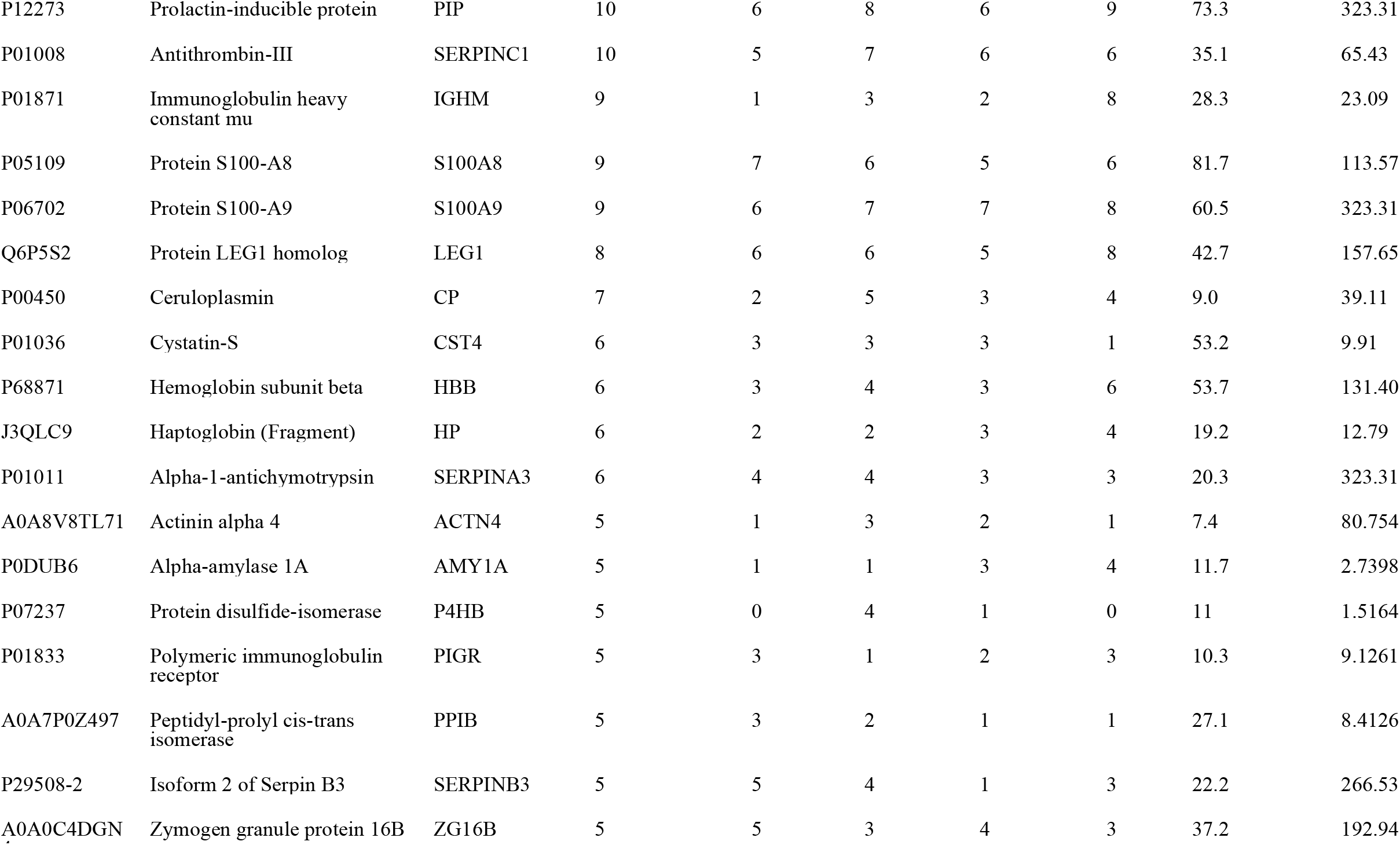
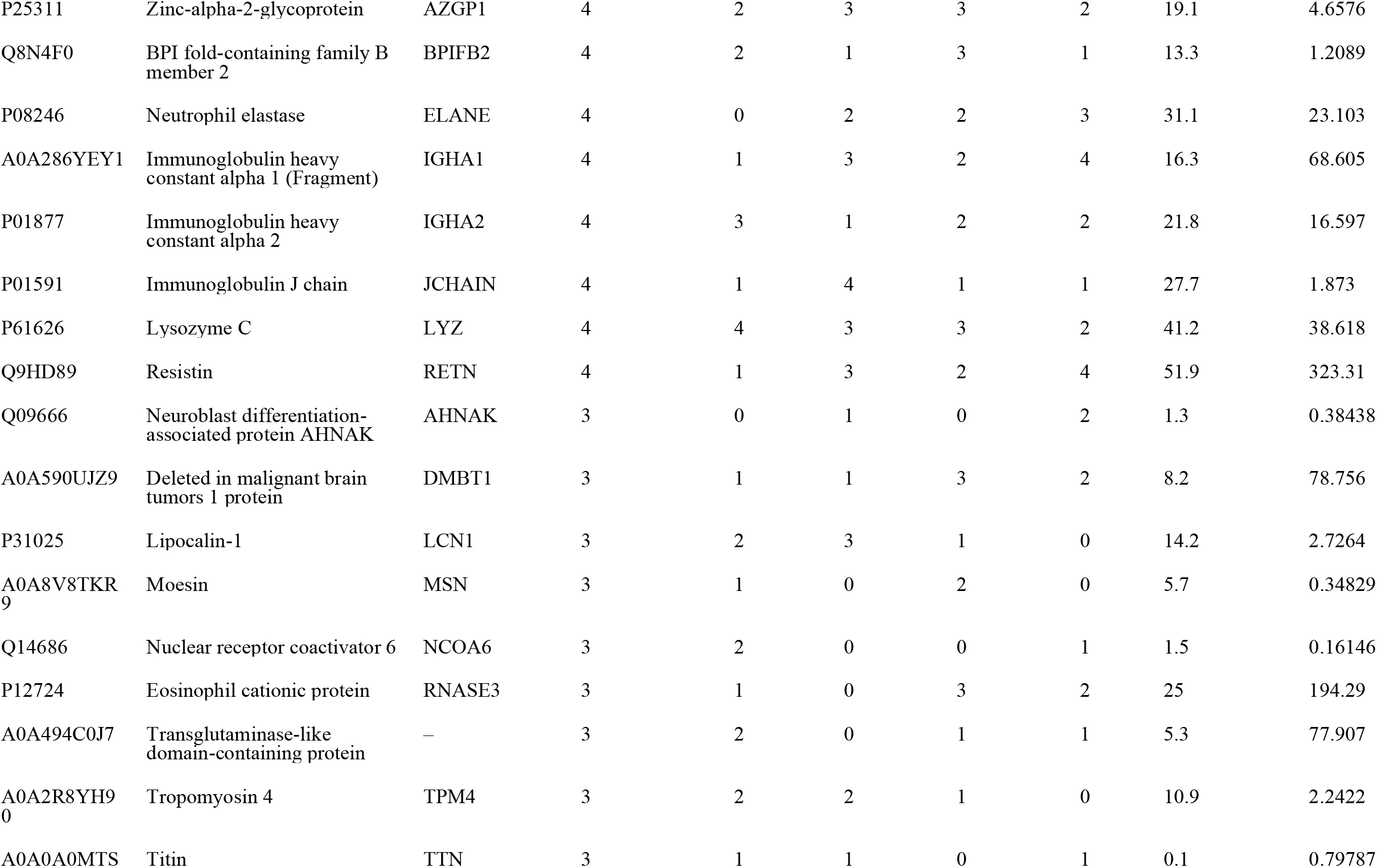
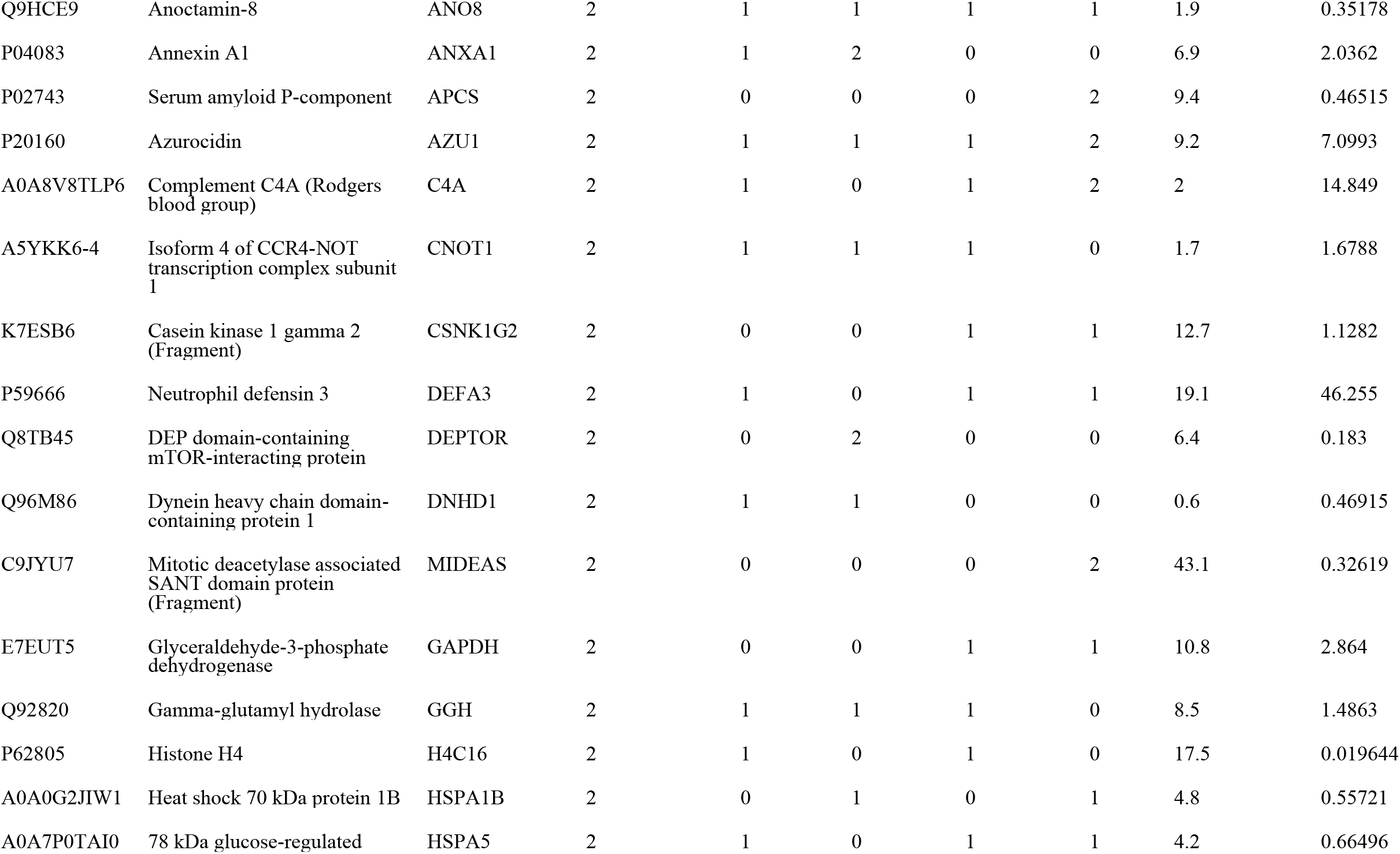
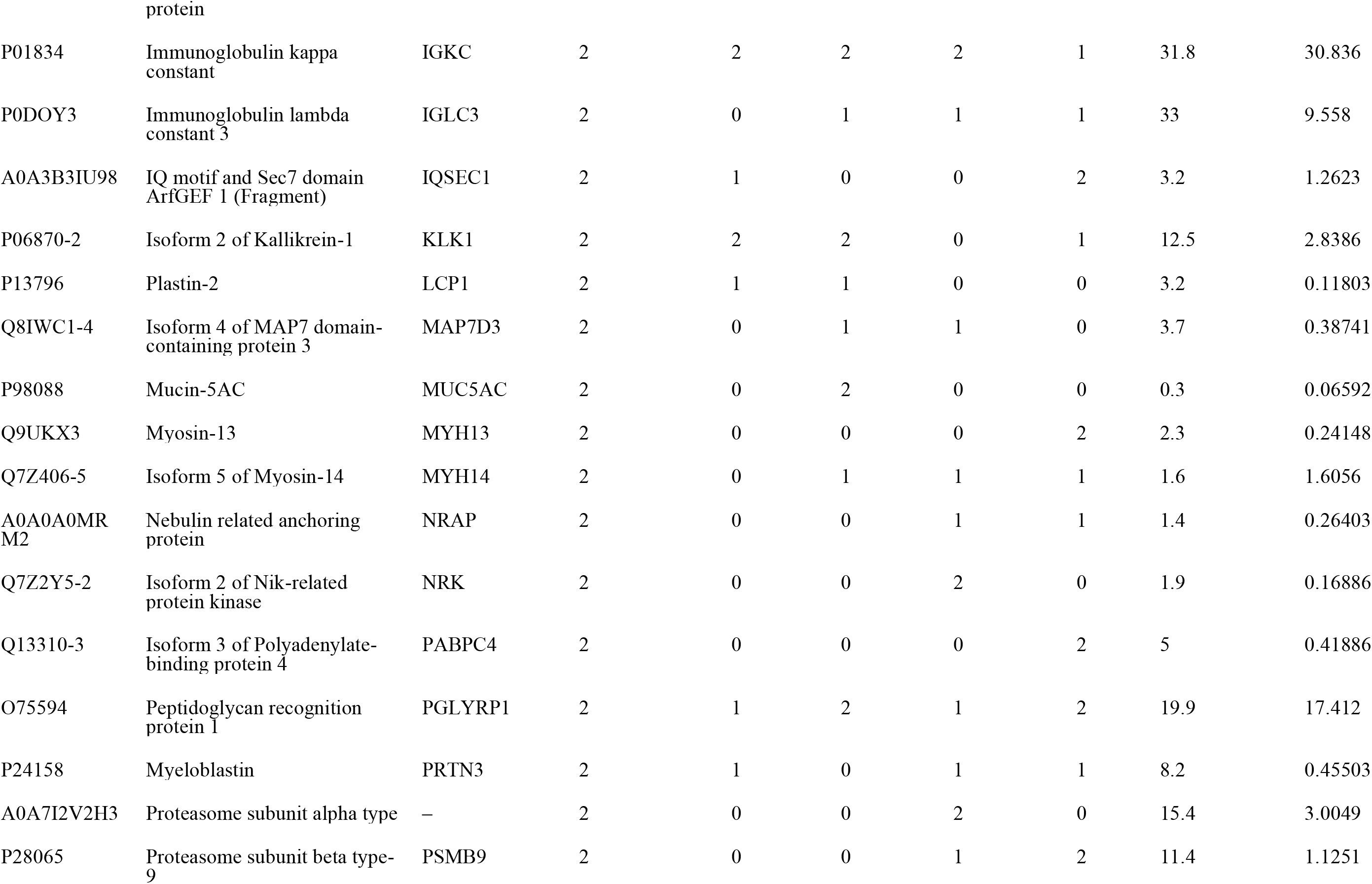
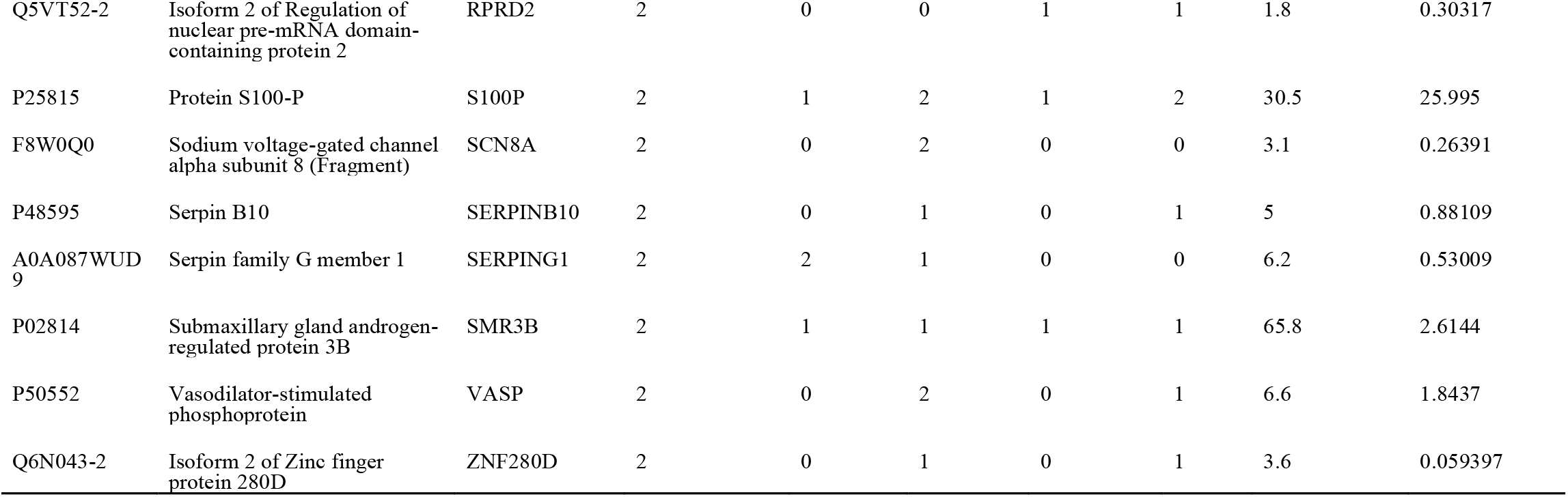
Human protein groups identified in the dental calculus of HM2-HA-3.

**Table 3.**
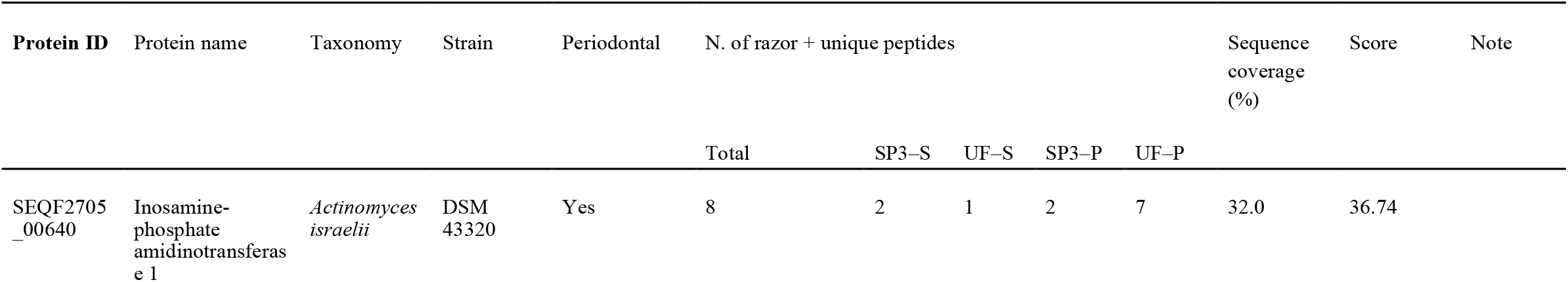
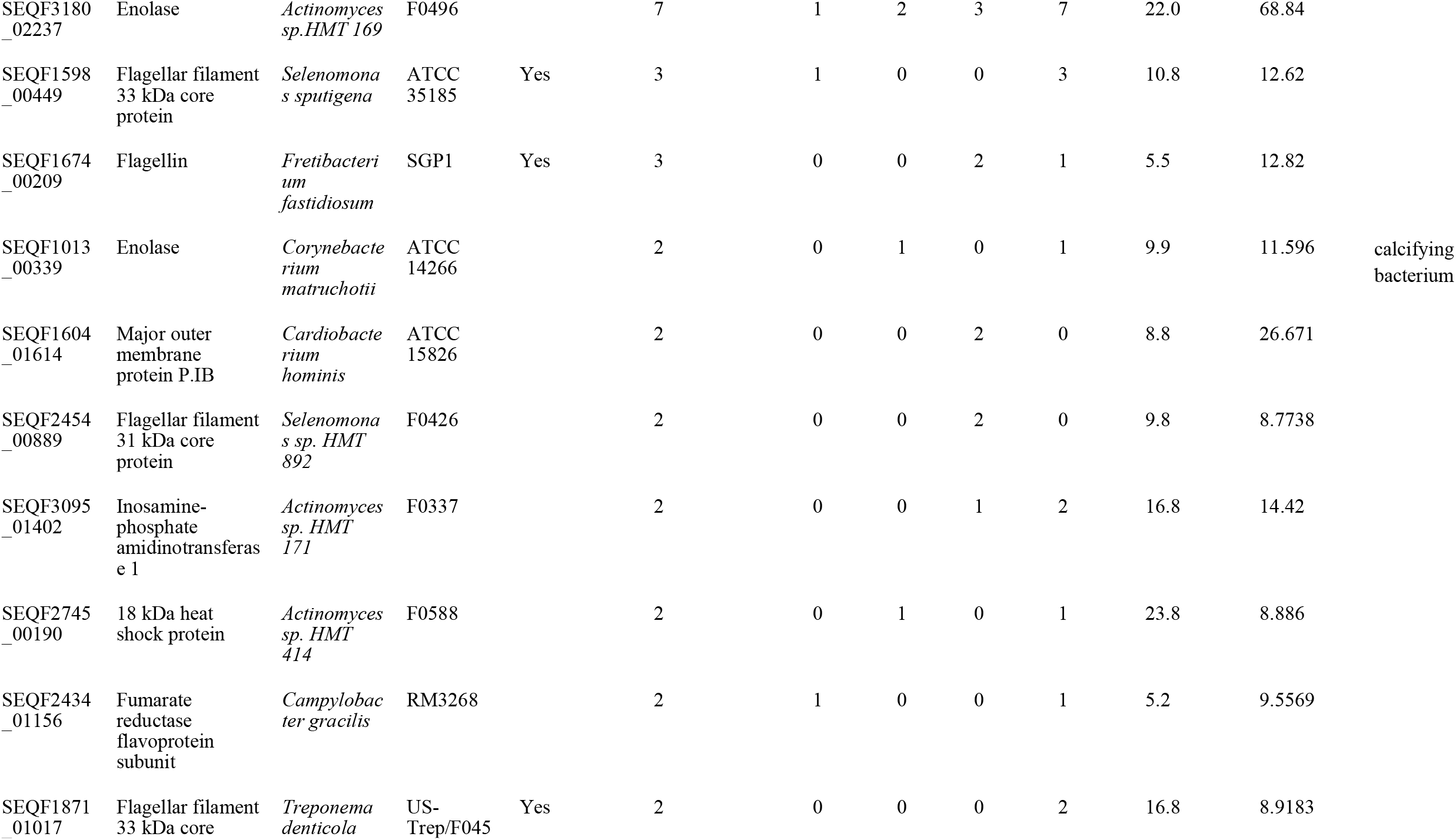
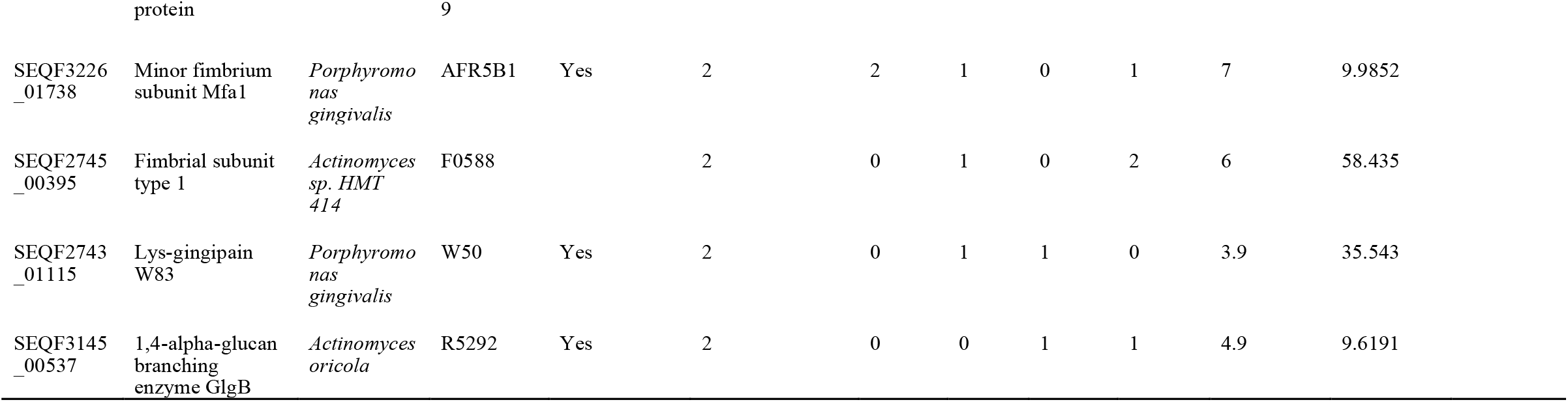
Oral bacterial protein groups identified in the dental calculus of HM2-HA-3.

The identified human proteins were classified with GO term using the PANTHER software^38^. Among the assigned protein class, 13.9% represented the “defense/immunity.” Among the proteins categorized in this class, peptidoglycan recognition protein 1 was one of the innate immune system proteins and functions to directly kill bacteria by recognizing and cleaving peptidoglycans on the bacterial wall^52^. Neutrophil elastase is among the antimicrobial peptides abundant in the saliva and gingival crevicular fluid in the oral cavity and is involved in local defense mechanisms^53^.

We identified a total of 15 proteins from 13 bacterial taxa from the calculus. Eight of these originated from six bacterial taxa that are reportedly associated with periodontal disease in modern patients (Table 3). We identified two of the three “red complex” bacteria, the most notable core bacterial species in the severe form of periodontal disease (*Porphyromonas gingivalis* and *Treponema denticola*). In addition, among the identified bacterial taxa, *Selenomonas sputigena* and *Fretibacterium fastidiosum* are reportedly associated with severe periodontal disease in modern humans^54,55^, while *Actinomyces dentalis* and *Actinomyces israelii* were identified in patients with severe periodontal disease^56^. *P. gingivalis* toxin, a proteolytic enzyme of Lys-gingipain W83, was identified in the calculus with well-annotated MS2 spectra (Supplementary Figure S2)^57^. Moreover, pathologically invasive proteins, such as *T. denticola* flagellar filament 33-kDa core protein, *F. fastidiosum* flagellin, and *S. sputigena* flagellar filament 33-kDa core protein, were also identified with well-annotated MS2 spectra (Supplementary Figure S2). These flagellar proteins are associated with bacterial motility and could initiate immune responses by interacting with toll-like receptor 5 in the host^58–61^. We could not identify any bacterial taxa and dental caries-associated proteins. Our BLAST search indicated that the peptide sequences of these periodontal disease-associated bacterial proteins only occur in certain bacterial genera (Supplementary Table S2).

We compared the protein groups or bacterial taxa identified in the dental calculus of HM2-HA-3 with those identified in a previous palaeoproteomic analysis of ancient human dental calculus from medieval Dalheim, Germany as well as those of modern European patients with periodontitis and dental caries^12^. As presented in Figure 3, 49.4% (40/81) of the human proteins and 69.2% (9/15) of the bacterial taxa identified in HM2-HA-3 calculus were also identified either in Dalheim or modern calculus^12^. with the common bacterial taxa being *P. gingivalis, A. israelii, Actinomyces sp*. HMT 414, and *Corynebacterium matruchotii*. Bacterial species unique to HM2-HA-3 included *S. sputigena, Actinomyces sp*. HMT 169, *Selenomonas sp*. HMT 892, and *Campylobacter gracilis*^*62*^.

**Figure 3.**
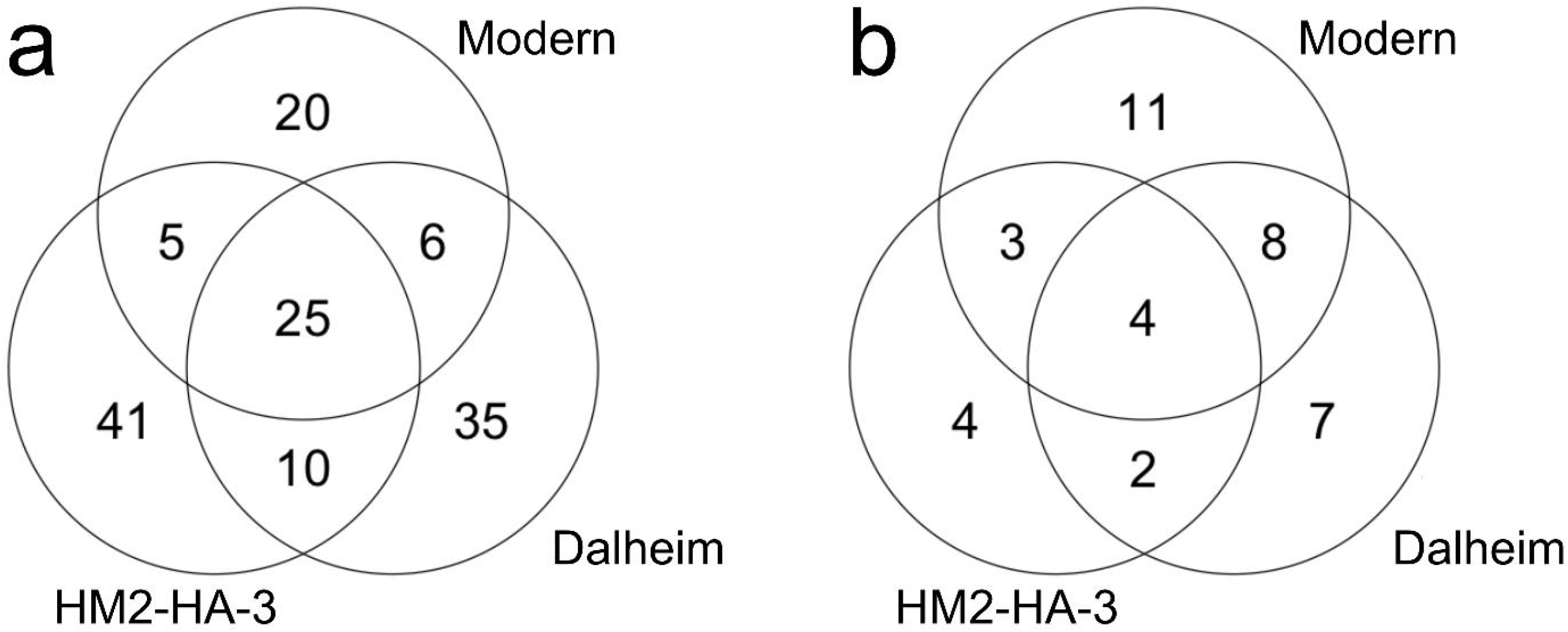
Venn diagrams of a) human proteins and b) bacterial taxa identified in the ancient dental calculus of HM2-HA-3 (this study) as well as in the dental calculus samples from medieval Dalheim and modern patients^12^.

The “defense/immunity” protein class proportion calculated by PANTHER was similar between the Dalheim (10.4%) and HM2-HA-3 (13.9%) calculi while that in modern calculus was higher (20.8%). The proportion of “immune system”-assigned biological processes calculated by PANTHER was lower in HM2-HA-3 (6.9%) than in Dalheim (8.1%) and modern (10.8%) dental calculi.

Finally, we performed a proteomic analysis of a rib bone sample of HM2-HA-3 to investigate the potential presence of systematic diseases. We identified a total of 59 human proteins, most of them being bone proteins (Supplementary Table S3). We could not identify any systematic disease-associated protein.

## Discussion

The palaeoproteomic analysis of abnormally deposited dental calculus conducted here provided molecular insights into the pathological conditions of the oral cavity of HM2-HA-3. We identified both pathogenic factors and bioinvasive proteins (i.e., Lys-gingipain W83, flagellin, and flagellar filament 33-kDa) from bacterial taxa reportedly associated with periodontal disease in modern patients. The identification of these proteins provides molecular support for the periodontal disease of this individual originally diagnosed based solely on physical characteristics. These bacterial proteins are associated with periodontal disease pathogenesis and development as well as with the secretion of inflammatory cytokines^58–61,63^. Of the 13 bacterial taxa identified from the calculus of HM2-HA-3, seven (53.8%) are reportedly associated with periodontal disease in modern clinical medicine (Table 3), in particular, two of the three red complex bacterial taxa. Proteins from the red complex bacteria have frequently been identified in both modern and ancient human dental calculus samples^12,14,64,65^. In this study, the pathogenic protein of *P. gingivalis* and bioinvasive protein of *T. denticola* were confidently identified^66^, providing direct evidence of red complex bacterial involvement in periodontal disease etiology. Although the involvement of the remaining seven bacterial taxa in the etiology of periodontal disease remains unclear, our results confidently indicate that periodontal disease bacterial etiology in HM2-HA-3 was similar to that in modern patients.

The presence of various host defense response proteins suggests that HM2-HA-3 was subjected to pathological stress and the resulting inflammation, at least during dental calculus deposition. However, the identified host defense proteins were nonspecific (e.g., lactotransferrin, immunoglobulin kappa constant, and prolactin-inducible protein) and mostly similar to those identified in other ancient individuals with significantly lower calculus deposition (Supplementary Figure S3)^12^. Moreover, our PANTHER analysis revealed that the “immune system process” comprised 6.9% of the total processes assigned to the identified host proteins in the HM2-HA-3 dental calculus (Figure 4). This proportion is rather lower compared to those in the calculus samples from medieval Dalheim (8.1%) and modern patients suffering from moderate to moderate/severe periodontal disease (10.7%)^12^. Furthermore, the proportion of the “defense/immunity protein” class was also lower in the calculus of HM2-HA-3 (13.9%) than that in the modern dental calculus (20.8%) and was somewhat higher than that in the calculus sample of medieval Dalheim (10.4%)^12^. These results imply that host defense response to oral pathological stress was not necessarily higher in HM2-HA-3, who exhibited significant amounts of calculus deposits and severe masticatory dysfunction, relative to modern periodontitis patients and medieval individuals with lower calculus deposition.

**Figure 4.**
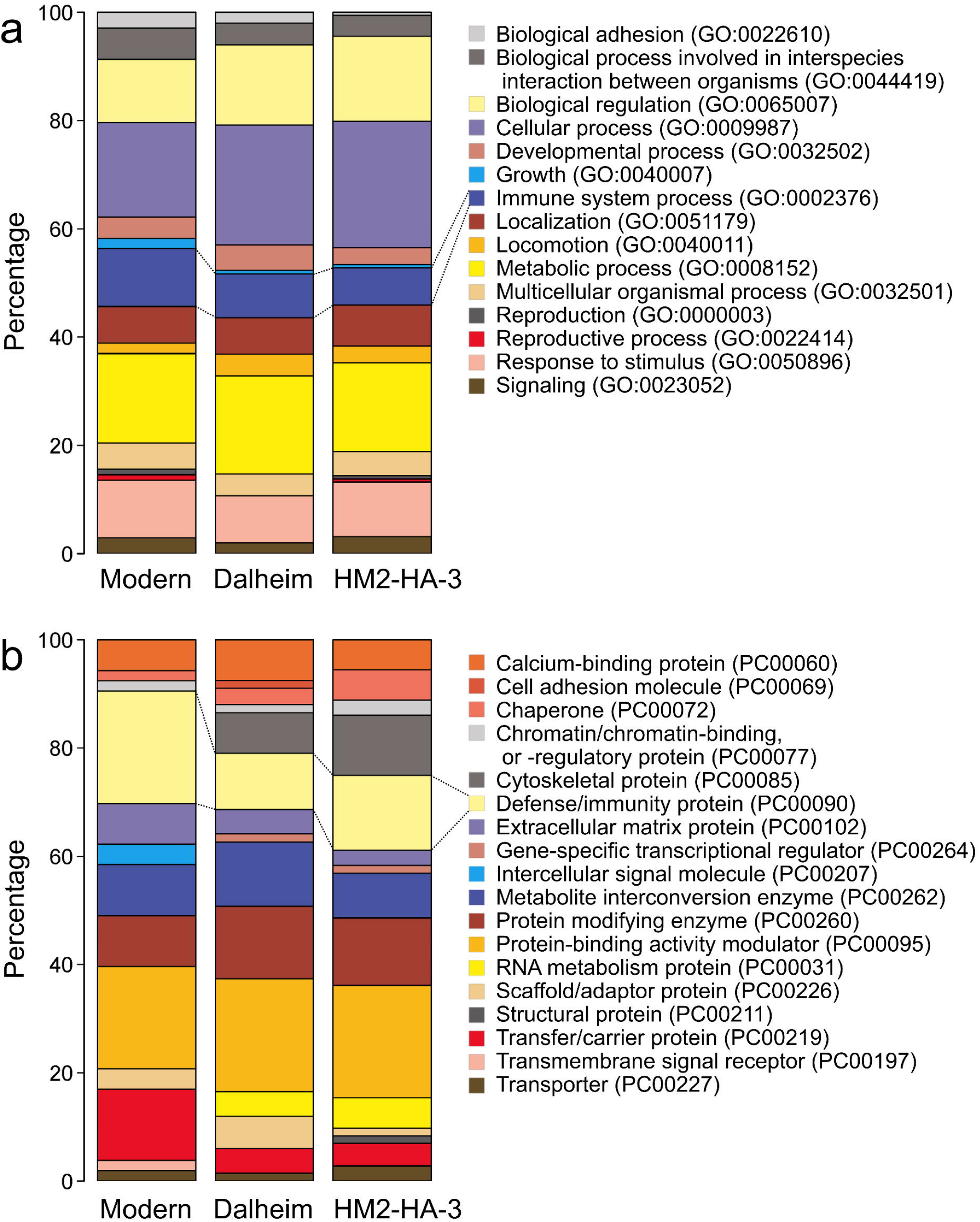
Results of PANTHER a) biological process and b) protein class analysis of protein groups identified in the dental calculus of HM2-HA-3 (this study) as well as in the dental calculus samples from medieval Dalheim and modern patients^12^.

Although palaeoproteomics provides molecular evidence on the bacterial etiology of and host defense response to periodontal disease, the cause of the abnormal calculus deposition in HM2-HA-3 remains unclear. Diet is often cited as a cause for calculus deposition^67^, but this cause is unlikely for HM2-HA-3. Stable isotope analysis showed that HM2-HA-3 had a similar diet to other individuals from the Hamanaka 2 site and other individuals from Hamanaka 2 site displayed little or no calculus deposition (Figure 2). Abnormally high amounts of calculus deposition could occasionally be seen in modern patients, but the underlying cause is unidentifiable in most cases^68,69^. At least, this individual would not have a routine tooth cleaning habit during the period of calculus deposition. Furthermore, as the HM2-HA-3 bone proteome did not contain disease-indicative proteins, calculus deposition unlikely occurred as a systemic disease byproduct.

HM2-HA-3 is the first individual among the ancient human skeletons from Asia with a bacterial proteome studied in detail. Therefore, in this study, we used for comparison previously published proteome results on calculi from individuals in Europe^12^. Almost all published bacterial proteome of modern and ancient dental calculus originate from Europe^12,14^. As the regional differences in the human oral bacterial composition have been suggested^70^, accumulating data on dental calculus bacterial proteome outside Europe would be required.

## Materials and Methods

Detailed procedures regarding sample collection and analyses are described in the Supplementary Information. A brief summary is shown below.

### Sampling

Dental calculus was collected from the lower right first incisor of HM2-HA-3 (Supplementary Figure S1), with the method described previously^29^. Given the small variability in bacterial composition in calculus obtained from different oral positions within an individual^30^, we assume that this sample had a representative bacterial composition as would be obtained from the abnormally deposited calculus present on the molars (Figure 1). Rib bones were also sampled for palaeoproteomic and isotope analyses.

### Proteomics

Protein extraction from 15 mg of dental calculus was performed using modified ultrafiltration and single-pot solid-phase-enhanced sample preparation (SP3) methods for ancient protein analysis^31,32^. Protein extraction from 20 mg rib bone was performed using modified ultrafiltration method^33^. Following the guidelines for palaeoproteomics^17^, the entire extraction process was carried out in a clean laboratory dedicated to ancient biomolecules built at the Graduate University for Advanced Studies, Japan. We obtained four fractions of the calculus sample (i.e., supernatant and pellet fractions from each of the ultrafiltration and SP3 methods) and two fractions (i.e., supernatant and pellet) of bone sample along with experimental blanks. Liquid chromatgraphy-tandem mass spectrometry (LC-MS/MS) analysis of dental calculus was performed using an Orbitrap Fusion Tribrid mass spectrometer (Thermo Fisher Scientific) at Japan Agency for Marine-Earth Science and Technology (JAMSTEC) with the conditions described in Nunoura et al.^34^. LC-MS/MS analysis of rib bone was performed using an Orbitrap QE Plus mass spectrometer (Thermo Fisher Scientific) at Kanazawa University with the conditions described in Ogura et al.^35^. RAW data files generated by LC-MS/MS were analyzed using the MaxQuant software version 2.0.1.0^36^. Data of calculus were searched against the Oral Signature Screening Database (OSSD^19^) for the first quality-assuarance step and the electric Human Oral Microbiome Database (eHOMD^37^) or entire human proteome (as of 2023-03-02) for the second protein identification step. Data of bone were searched against the entire human proteome. Because no food proteins was identified from dental calculus in a MaxQuant search against an entire Swiss-Prot database (as of 2021-08-20), we did not investigate into food proteins. Comparative datasets were analyzed anew in the same manner^12^.

Gene Ontology (GO) analysis of the human-derived proteins identified from the dental calculus of HM2-HA-3 was performed using PANTHER, version 14^38^. Python script reported by Mackie et al.^13^ was used to calculate asparagine and glutamine deamidation rates. All sunsequrnt data analyses were performed using R, version 4.2.2 (R Core Team, 2022).

### Radiocarbon dating and stable isotope analysis

Collagen was extracted from a rib bone of HM2-HA-3 to conduct radiocarbon measurement and carbon and nitrogen stable isotope analysis, based on the method described previously^39^. Carbon and nitrogen stable isotopes were measured using elemental analyzer-isotope ratio mass spectrometry (EA-IRMS) at the University Museum, the University of Tokyo (UMUT).

Radiocarbon concentrations were measured using accelerator mass spectrometry (AMS) at UMUT. Radiocarbon age was calibrated against atmospheric and marine calibration curves (IntCal20 and Marine20^40,41^) and with the local marine reservoir effect^42^ using OxCal, version 4.4^43^.

## Supporting information

Supplemental imformation of Palaeoproteomic investigation of an ancient human skeleton with abnormal deposition of dental calculus

## Acknowledgments

This study was supported in part by Grants-in-Aid for Scientific Research (KAKENHI: 16H06408, 19K01134, 19K06868, 20H01370, 20H05821, 20KK0166, 21H00588, 22KK0170, 23H00032) from Japan Society for the Promotion of Science. The authors thank Ken Takai for providing supports to this study.

## Author contribution

Conceptualization: YU-F, RS, TT; Investigation: YU-F, SS, RS, TN, MY, TT; Resources: IH, HM; Writing - Original Draft: YU-F, TT; Visualization: YU-F, TT

## Competing interests

The authors declare no competing interests.

## Data availability

LC-MS/MS data have been uploaded to PRIDE repository^71^ with the dataset identifier PXD044070.

## Notes

### Competing Interest Statement

The authors have declared no competing interest.

## References

1. Kilgore, L., Jurmain, R. & Van Gerven, D. Palaeoepidemiological patterns of trauma in a medieval Nubian skeletal population. Int. J. Osteoarchaeol. 7, 103–114 (1997).

2. Betsinger, T. K. & Smith, M. O. A singular case of advanced caries sicca in a pre-Columbian skull from East Tennessee. Int J Paleopathol 24, 245–251 (2019).

3. Tilley, L. & Oxenham, M. F. Survival against the odds: Modeling the social implications of care provision to seriously disabled individuals. Int J Paleopathol 1, 35–42 (2011).

4. Milella, M., Zollikofer, C. P. E. & Ponce de León, M. S. Virtual reconstruction and geometric morphometrics as tools for paleopathology: a new approach to study rare developmental disorders of the skeleton. Anat. Rec. 298, 335–345 (2015).

5. Majander, K. et al. Ancient Bacterial Genomes Reveal a High Diversity of Treponema pallidum Strains in Early Modern Europe. Curr. Biol. 30, 3788–3803.e10 (2020).

6. Andrades Valtueña, A. et al. Stone Age Yersinia pestis genomes shed light on the early evolution, diversity, and ecology of plague. Proc. Natl. Acad. Sci. U. S. A. 119, e2116722119 (2022).

7. Akcalı, A. & Lang, N. P. Dental calculus: the calcified biofilm and its role in disease development. Periodontol. 2000 76, 109–115 (2018).

8. Brothwell, D. R. Digging Up Bones: The Excavation, Treatment, and Study of Human Skeletal Remains. (Cornell University Press, 1981).

9. Hanihara, T., Ishida, H., Ohshima, N., Kondo, O. & Masuda, T. Dental calculus and other dental disease in a human skeleton of the Okhotsk Culture unearthed at Hamanaka-2 site, Rebun-Island, Hokkaido, Japan. Int. J. Osteoarchaeol. 4, 343–351 (1994).

10. Wright, S. L., Dobney, K. & Weyrich, L. S. Advancing and refining archaeological dental calculus research using multiomic frameworks. STAR: Science & Technology of Archaeological Research 7, 13–30 (2021).

11. Radini, A. & Nikita, E. Beyond dirty teeth: Integrating dental calculus studies with osteoarchaeological parameters. Quaternary International Preprint at https://doi.org/10.1016/j.quaint.2022.03.003 (2022).

12. Warinner, C. et al. Pathogens and host immunity in the ancient human oral cavity. Nat. Genet. 46, 336–344 (2014).

13. Mackie, M. et al. Palaeoproteomic Profiling of Conservation Layers on a 14th Century Italian Wall Painting. Angew. Chem. Int. Ed Engl. 57, 7369–7374 (2018).

14. Jersie-Christensen, R. R. et al. Quantitative metaproteomics of medieval dental calculus reveals individual oral health status. Nat. Commun. 9, 4744 (2018).

15. Fotakis, A. K. et al. Multi-omic detection of Mycobacterium leprae in archaeological human dental calculus. Philos. Trans. R. Soc. Lond. B Biol. Sci. 375, 20190584 (2020).

16. Warinner, C. et al. Direct evidence of milk consumption from ancient human dental calculus. Sci. Rep. 4, 7104 (2014).

17. Hendy, J. et al. A guide to ancient protein studies. Nature Ecology & Evolution 2, 791–799 (2018).

18. Jeong, C. et al. Bronze Age population dynamics and the rise of dairy pastoralism on the eastern Eurasian steppe. Proc. Natl. Acad. Sci. U. S. A. 115, E11248–E11255 (2018).

19. Wilkin, S. et al. Dairying enabled Early Bronze Age Yamnaya steppe expansions. Nature 598, 629–633 (2021).

20. Scorrano, G. et al. Genomic ancestry, diet and microbiomes of Upper Palaeolithic hunter-gatherers from San Teodoro cave. Commun Biol 5, 1262 (2022).

21. Tang, L. et al. Paleoproteomic evidence reveals dairying supported prehistoric occupation of the highland Tibetan Plateau. Sci Adv 9, eadf0345 (2023).

22. Ishida, H., Haniwara, T. & Kondo, O. [Human remains from the Hamanaka 2 Site, Rebun Island. ] Rebunto hamanaka 2 iseki shutsudo no jinkotsu (daini/yon ji chosa) (in Japanese). Tsukuba archaeological studies/ University of Tsukuba Prehistory and Archaeology Editorial Board, ed, 89–108 (2002).

23. Hudson, M. J. The perverse realities of change: world system incorporation and the Okhotsk culture of Hokkaido. Journal of Anthropological Archaeology 23, 290–308 (2004).

24. Naito, Y. I. et al. Dietary Reconstruction of the Okhotsk Culture of Hokkaido, Japan, Based on Nitrogen Composition of Amino Acids: Implications for Correction of 14C Marine Reservoir Effects on Human Bones. Radiocarbon 52, 671–681 (2010).

25. Tsutaya, T., Naito, Y. I., Ishida, H. & Yoneda, M. Carbon and nitrogen isotope analyses of human and dog diet in the Okhotsk culture: perspectives from the Moyoro site, Japan. Anthropol. Sci. 122, 89–99 (2014).

26. Leipe, C. et al. Barley (Hordeum vulgare) in the Okhotsk culture (5th–10th century AD) of northern Japan and the role of cultivated plants in hunter–gatherer economies. PLoS One 12, e0174397 (2017).

27. Ohshima, N. [ Historical trends in caries frequency in archaeological human bones from Hokkaido, Japan] Hokkaido no kojinkotsu ni okeru ushokuhindo no jidaitekisuii(in Japanese). Anthropol. Sci. 104, 385–397 (1996).

28. Oxenham, M. F. & Matsumura, H. Oral and physiological paleohealth in cold adapted peoples: Northeast Asia, Hokkaido. Am. J. Phys. Anthropol. 135, 64–74 (2008).

29. Sawafuji, R., Saso, A., Suda, W., Hattori, M. & Ueda, S. Ancient DNA analysis of food remains in human dental calculus from the Edo period, Japan. PLoS One 15, e0226654 (2020).

30. Fagernäs, Z. et al. Understanding the microbial biogeography of ancient human dentitions to guide study design and interpretation. FEMS Microbes 3, xtac006 (2022).

31. Hughes, C. S. et al. Single-pot, solid-phase-enhanced sample preparation for proteomics experiments. Nat. Protoc. 14, 68–85 (2019).

32. Palmer, K. S. et al. Comparing the Use of Magnetic Beads with Ultrafiltration for Ancient Dental Calculus Proteomics. J. Proteome Res. 20, 1689–1704 (2021).

33. Sawafuji, R. et al. Proteomic profiling of archaeological human bone. R Soc Open Sci 4, 161004 (2017).

34. Nunoura, T. et al. A primordial and reversible TCA cycle in a facultatively chemolithoautotrophic thermophile. Science 359, 559–563 (2018).

35. Ogura, K. et al. Potential biomarker proteins for aspiration pneumonia detected by shotgun proteomics using buccal mucosa samples: a cross-sectional case–control study. Clin. Proteomics 20, 9 (2023).

36. Tyanova, S., Temu, T. & Cox, J. The MaxQuant computational platform for mass spectrometry-based shotgun proteomics. Nat. Protoc. 11, 2301–2319 (2016).

37. Chen, T. et al. The Human Oral Microbiome Database: a web accessible resource for investigating oral microbe taxonomic and genomic information. Database 2010, baq013 (2010).

38. Mi, H., Muruganujan, A., Ebert, D., Huang, X. & Thomas, P. D. PANTHER version 14: more genomes, a new PANTHER GO-slim and improvements in enrichment analysis tools. Nucleic Acids Res. 47, D419–D426 (2019).

39. Tsutaya, T., Gakuhari, T., Asahara, A. & Yoneda, M. Isotopic comparison of gelatin extracted from bone powder with that from bone chunk and development of a framework for comparison of different extraction methods. Journal of Archaeological Science: Reports 11, 99–105 (2017).

40. Heaton, T. J. et al. Marine20—The Marine Radiocarbon Age Calibration Curve (0–55,000 cal BP). Radiocarbon 62, 779–820 (2020).

41. Reimer, P. J. et al. The IntCal20 Northern Hemisphere Radiocarbon Age Calibration Curve (0–55 cal kBP). Radiocarbon 62, 725–757 (2020).

42. Yoneda, M. et al. Radiocarbon marine reservoir ages in the western Pacific estimated by pre-bomb molluscan shells. Nucl. Instrum. Methods Phys. Res. B 259, 432–437 (2007).

43. Ramsey, C. B. Radiocarbon Calibration and Analysis of Stratigraphy: The OxCal Program. Radiocarbon 37, 425–430 (1995).

44. DeNiro, M. J. Postmortem preservation and alteration of in vivo bone collagen isotope ratios in relation to palaeodietary reconstruction. Nature 317, 806–809 (1985).

45. van Klinken, G. J. Bone Collagen Quality Indicators for Palaeodietary and Radiocarbon Measurements. J. Archaeol. Sci. 26, 687–695 (1999).

46. Junno, A. et al. Building a high-resolution chronology for northern Hokkaido – A case study of the Late Holocene Hamanaka 2 site on Rebun Island, Hokkaido (Japan). J. Archaeol. Sci. Rep. 36, 102867 (2021).

47. Howland, M. R. et al. Expression of the dietary isotope signal in the compound-specific ?13C values of pig bone lipids and amino acids. Int. J. Osteoarchaeol. 13, 54–65 (2003).

48. Hedges, R. E. M., Clement, J. G., Thomas, C. D. L. & O’connell, T. C. Collagen turnover in the adult femoral mid-shaft: modeled from anthropogenic radiocarbon tracer measurements. Am. J. Phys. Anthropol. 133, 808–816 (2007).

49. Okamoto, Y. et al. An Okhotsk adult female human skeleton (11th/12th century AD) with possible SAPHO syndrome from Hamanaka 2 site, Rebun Island, northern Japan. Anthropol. Sci. 124, 107–115 (2016).

50. van Doorn, N. L., Wilson, J., Hollund, H., Soressi, M. & Collins, M. J. Site-specific deamidation of glutamine: a new marker of bone collagen deterioration. Rapid Commun. Mass Spectrom. 26, 2319–2327 (2012).

51. Wilson, J., van Doorn, N. L. & Collins, M. J. Assessing the extent of bone degradation using glutamine deamidation in collagen. Anal. Chem. 84, 9041–9048 (2012).

52. Kashyap, D. R. et al. Peptidoglycan recognition proteins kill bacteria by inducing oxidative, thiol, and metal stress. PLoS Pathog. 10, e1004280 (2014).

53. Dolińska, E. et al. The Effect of Nonsurgical Periodontal Therapy on HNP1-3 Level in Gingival Crevicular Fluid of Chronic Periodontitis Patients. Arch. Immunol. Ther. Exp. 65, 355–361 (2017).

54. Oliveira, R. R. D. S. et al. Levels of Candidate Periodontal Pathogens in Subgingival Biofilm. J. Dent. Res. 95, 711–718 (2016).

55. Curtis, M. A., Diaz, P. I. & Van Dyke, T. E. The role of the microbiota in periodontal disease. Periodontol. 2000 83, 14–25 (2020).

56. Vielkind, P., Jentsch, H., Eschrich, K., Rodloff, A. C. & Stingu, C.-S. Prevalence of Actinomyces spp. in patients with chronic periodontitis. Int. J. Med. Microbiol. 305, 682–688 (2015).

57. Genco, C. A., Potempa, J., Mikolajczyk-Pawlinska, J. & Travis, J. Role of gingipains R in the pathogenesis of Porphyromonas gingivalis-mediated periodontal disease. Clin. Infect. Dis. 28, 456–465 (1999).

58. Hayashi, F. et al. The innate immune response to bacterial flagellin is mediated by Toll-like receptor 5. Nature 410, 1099–1103 (2001).

59. Mahanonda, R. & Pichyangkul, S. Toll-like receptors and their role in periodontal health and disease. Periodontol. 2000 43, 41–55 (2007).

60. Kim, C. et al. Immunotherapy targeting toll-like receptor 2 alleviates neurodegeneration in models of synucleinopathy by modulating α-synuclein transmission and neuroinflammation. Mol. Neurodegener. 13, 43 (2018).

61. Rath, C. B. et al. Flagellin Glycoproteomics of the Periodontitis Associated Pathogen Selenomonas sputigena Reveals Previously Not Described O-glycans and Rhamnose Fragment Rearrangement Occurring on the Glycopeptides. Mol. Cell. Proteomics 17, 721–736 (2018).

62. Tanner, A., Maiden, M. F., Macuch, P. J., Murray, L. L. & Kent, R. L., Jr. Microbiota of health, gingivitis, and initial periodontitis. J. Clin. Periodontol. 25, 85–98 (1998).

63. Kopeckova, M., Pavkova, I. & Stulik, J. Diverse Localization and Protein Binding Abilities of Glyceraldehyde-3-Phosphate Dehydrogenase in Pathogenic Bacteria: The Key to its Multifunctionality? Front. Cell. Infect. Microbiol. 10, 89 (2020).

64. Mackie, M. et al. Preservation of the metaproteome: variability of protein preservation in ancient dental calculus. Sci Technol Archaeol Res 3, 74–86 (2017).

65. Velsko, I. M. et al. Microbial differences between dental plaque and historic dental calculus are related to oral biofilm maturation stage. Microbiome 7, 102 (2019).

66. Socransky, S. S. & Haffajee, A. D. Periodontal microbial ecology. Periodontol. 2000 38, 135–187 (2005).

67. Lieverse, A. R. Diet and the aetiology of dental calculus. International Journal of Osteoarchaeology vol. 9 219–232 Preprint at https://doi.org/10.1002/(sici)1099-1212(199907/08)9:4<219::aid-oa475>3.0.co;2-v (1999).

68. Miyake, M., Iwasaki, A., Saito, H., Ohbayashi, Y. & Nagahata, S. A case of a giant dental calculus suspected to be a neoplastic lesion. Jpn. J. Oral Maxillofac. Surg. 50, 442–445 (2004).

69. Iwama, R. et al. A case of giant dental calculus in a patient with centronuclear myopathy. Spec. Care Dentist. (2022) doi:10.1111/scd.12772.

70. Gupta, V. K., Paul, S. & Dutta, C. Geography, Ethnicity or Subsistence-Specific Variations in Human Microbiome Composition and Diversity. Front. Microbiol. 8, 1162 (2017).

71. Perez-Riverol, Y. et al. The PRIDE database resources in 2022: a hub for mass spectrometry-based proteomics evidences. Nucleic Acids Res. 50, D543–D552 (2022).

