## Supplemental imformation of Palaeoproteomic investigation of an ancient human skeleton with abnormal deposition of dental calculus for "Palaeoproteomic investigation of an ancient human skeleton with abnormal deposition of dental calculus"

### Supplementary Materials and Methods

*Sampling*

HM2-HA-3 is housed in the School of Health Science, Sapporo Medical University. The sampling was performed as described previously (Sawafuji et al., 2020). Supragingival calculus was collected from the buccal surface of the lower right permanent first incisor using a sterilized dental explorer (YDM Corporation). Considering the small variability of bacterial composition in calculus obtained from different positions within an individual (Fagernäs et al., 2022), we assumed this sample represents the entire calculus. The dental calculus was collected into a 1.5 mL DNA LoBind tube (Eppendorf). Masks, nitrile gloves, hairnets, and laboratory coats were worn throughout the process.

*Proteomics of dental calculus*

Protein extraction from dental calculus was performed using modified ultrafiltration and paramagnetic bead methods for ancient proteins (Palmer et al., 2021). Following the guidelines for ancient protein research (Hendy et al., 2018), the entire process was carried out in a clean laboratory dedicated to ancient biomolecules built at the Graduate University for Advanced Studies. First, 15 mg of dental calculus was washed with 0.5 M EDTA for 5 minutes. Then, the calculus was smashed with micropestle and demineralized with 250 μL of 0.5 M EDTA for 3 days. The sample was centrifuged at 8000 ×g for 2 min and the supernatant was collected. The supernatant was reduced and alkylated with 10 mM TCEP and 20 mM CAA (Gu buffer) at 80 °C for 2 h under agitation. The pellet was denatured, reduced, and alkylated with 250 μL of a solution containing 2 M guanidine, 0.1 M Tris, 10 mM TCEP, and 20 mM CAA at 80 °C for 2 h under agitation. Protein LoBind tubes (Eppendorf) were used for the experimental process.

The supernatant and pellet solutions were divided into two aliquots for ultrafiltration and single-pot solid-phase-enhanced sample preparation (SP3) (Hughes et al., 2018). Each 100 μL of supernatant and pellet samples was loaded on Amicon 3000 Da molecular weight cut-off spin columns and centrifuged to exchange the buffer. The ultrafiltered samples were washed and resuspended with 0.1 M triethylammonium bicarbonate (TEAB) solution in new tubes. In parallel, each 100 μL of supernatant and pellet samples were mixed with a mixture of two hydrophilic paramagnetic beads (Sera-Mag SpeedBeads, GE Healthcare) at the final concentration of ~2 μg/μL. Then, 108 μL of 100% ethanol was added, the solution was gently mixed, and beads were aggregated with a magnet. The supernatant was removed, and the beads were washed with 180 μL of 80% ethanol three times. After washing, beads were aggregated with a magnet, ethanol was carefully removed, and TEAB solution was added. Samples suspended in TEAB solutions were digested with a total of 0.4 μg trypsin at 37℃ for 2 h and then 37℃ overnight. The digested samples were collected and acidified with TFA. The peptides were purified with spin columns filled with C18 resins, eluted with 70% ACN, and dried with a vacuum centrifuge. LC-MS/MS analysis was performed with the condition described in Nunoura et al. (2018). The dried peptides were resuspended in 2% acetonitrile and 0.1% trifluoroacetic acid in water. Peptides were separated using an Ultimate 3000 RSLCnano system (Thermo Fisher Scientific) with a reverse-phase Zaplous alpha Pep-C18 column (3 μm, 120 Å, 0.1 × 150 mm, AMR). The gradient was composed of solvent A (0.1% formic acid in water) and solvent B (100% acetonitrile). At the time of injection, the mobile phase was 5% solvent B at a flow rate of 300 nL per minute. The concentration of solvent B was linearly increased to 45% over 100 min. The column temperature was set at 35 °C via a nano-electrospray ion source (Dream Spray, AMR). The Orbitrap Fusion Tribrid mass spectrometer (Thermo Fisher Scientific) was operated in the positive ion mode. For peptide ionization, an electrospray voltage of 1.7 kV was applied, and the ion transfer tube temperature was set at 250 °C. All MS spectra were obtained in the Orbitrap mass analyzer ($m/z$ range was 350–1800 and the resolution was 120,000 full width at half maximum) with EASY-IC internal mass calibration. MS/MS spectra resulting from collision-induced dissociation (CID) fragmentation were obtained in an ion trap mass analyzer (*m/z* range was auto and scan rate was "rapid").

RAW data files generated by LC-MS/MS were analyzed using the MaxQuant software version 2.0.1.0 (Tyanova et al., 2016). In the first search, Oral Signature Screening Database (OSSD) was used to diagnose the authenticity of the proteome (Wilkin et al., 2021). In the second search, RAW data were searched against the electric Human Oral Microbiome Database (eHOMD) (Chen et al., 2010) or the entire human proteome (as of 2023-03-02). The protein groups that were identified with non-overlapping ≧2 unique and razor peptides in the second search were considered to be present in the samples. The following parameters were used for the analysis. Parent mass error and fragment mass tolerances were those pre-set for Orbitraps. Carbamidomethylation was set as a fixed modification, and oxidation of methionine, deamidation of Asparagine and Glutamine, hydroxyproline, and derivation of pyroglutamic acid were set at variable modifications. Up to a maximum of 5 modifications per peptide was allowed. All peptides were automatically filtered by a false discovery rate (FDR) of 1.0%. All contaminant accessions (i.e., keratins and trypsin) were excluded from further analysis using the contamination.fasta provided by MaxQuant, which includes common laboratory contaminants.

Gene Oncology (GO) analysis of the human-derived proteins identified from the dental calculus of HM2-HA-3 was performed using PANTHER, version 14 (Mi et al., 2019). Python script reported by Mackie et al. (2018) was used to calculate asparagine and glutamine deamidation rates.

*Proteomics of bone*

Protein extraction from a rib was performed using a modified ultrafiltration method for ancient proteins (Sawafuji et al., 2017). Following the guidelines for ancient protein research (Hendy et al., 2018), the entire process was carried out in a clean laboratory dedicated to ancient biomolecules built at the Graduate University for Advanced Studies. First, 20 mg of bone was demineralized with 500 µL of 0.5 M EDTA for 2 days. The sample was centrifuged at 8000 ×g for 2 min and the supernatant was collected. The supernatant was purified using Amicon Ultra Centrifugal Filters (molecular weight cutoff of 3000 Da) and resuspended in Gu buffer. The pellet was suspended in the Gu buffer. Both supernatant and pellet fractions were heated at 80 °C for 2 h under agitation to denature, reduce, and alkylate proteins. The concentrations of resultant protein solutions were measured by BCA assay. Then, the protein solutions were diluted with 25 mM Tris in 10% ACN to the final concentration of 0.6 M GuHCl. Protein solutions were digested using 0.4 µg Lys-C and 0.4 µg trypsin mix (Promega) overnight at 37°C. The digested samples were acidified with TFA. The peptides were purified with spin columns filled with C18 resins, eluted with 70% ACN, and dried with a vacuum centrifuge.

LC-MS/MS analysis was performed with the condition described in Ogura et al. (2023). The dried peptides were resuspended with solvent and loaded onto the nano-liquid chromatography EASY-nLC 1200 system (ThermoFisher Scientific). This system is equipped with a pre-column (Acclaim PepMap100 C18 column: inner diameter, 75 µm, length, 20 mm, particle size, 3.0 µm; ThermoFisher Scientific) and analytical column (Acclaim PepMap100 C18 column: inner diameter, 75 µm; length, 150 mm, particle size, 3.0 µm; ThermoFisher Scientific) equilibrated with 0.1% formic acid. Peptide elution is performed using a linear gradient (0–35%) of acetonitrile at a flow rate of 300 nL/min. The eluted peptides were ionized with a spray voltage of 2 kV (ion transfer tube temperature, 275 °C) and detected using LC-MS/MS (Thermo Orbitrap QE plus, ThermoFisher Scientific) in the data-dependent acquisition mode using Xcalibur (version 4.0; Thermo Fisher Scientific). Mass spectra with 375–1500 m/z were obtained with a resolution of 70,000 full width at half maximum.

RAW data files generated by LC-MS/MS were analyzed against the entire human proteome (as of 2023-03-02) using the MaxQuant software version 2.0.1.0 (Tyanova et al., 2016). The identification criteria and search parameters were the same as the analysis of dental calculus.

*Radiocarbon dating and stable isotope analysis*

Collagen was extracted from a rib bone of HM2-HA-3 to conduct radiocarbon measurement and carbon and nitrogen stable isotope analysis, based on the method described previously (Tsutaya et al., 2017). Briefly, the bone sample was mechanically cleaned and then sonicated in Milli-Q water to remove surface debris. The bone sample was then washed with 0.2 M NaOH overnight to remove humic substances. Then, the powdered bone sample was decalcified with 1 N HCl overnight at 4°C. The decalcified sample was concentrated, heated overnight at 90°C, and filtered by using a glass fiber filter (Whatman, GF/F). The filtered sample was freeze-dried and the resultant gelatin was used for the analyses.

Approximately 2.5 mg of extracted gelatin was burned into carbon dioxide, purified, and converted into graphite with a catalyst of iron powder. Radiocarbon concentration was measured using an accelerator mass spectrometry at the University Museum, the University of Tokyo (UMUT).

Radiocarbon age was calibrated against atmospheric and marine calibration curves (IntCal20 and Marine20, Reimer et al., 2020; Heaton et al., 2020), using the software OxCal, version 4.4 (Bronk Ramsey, 1995). Considering the marine-based diet of Okhotsk people (Naito et al., 2010; Tsutaya et al., 2014), the contribution from marine carbon was set as 80 ± 0%. The local marine reservoir effect (⊿R = -64 ± 36, recalculated with Marine20), measured for Soya Current affecting Rebun Island, was used for correction (Yoneda et al., 2007).

Approximately 35 mg of the extracted gelatin was loaded into a tin capsule and measured for carbon and nitrogen stable isotopes by an elemental analyzer-isotope ratio mass spectrometry (EA-IRMS; Thermo Flash 2000 elemental analyzer, Finnigan ConFlo IV interface, and Thermo Delta V mass spectrometer) at UMUT. Stable isotope ratios were calibrated by laboratory working standards (L-alanine 1: $\delta^{13}C = -18.5 \pm 0.2‰$, $\delta^{15}N = -1.02 \pm 0.2‰$; L-alanine 2: $\delta^{13}C = -19.6 \pm 0.2‰$, $\delta^{15}N = 8.7 \pm 0.2‰$; L- alanine 3: $\delta^{13}C = -19.6 \pm 0.2‰$; $\delta^{15}N = 20.0 \pm 0.2‰$; L-histidine: $\delta^{13}C = -7.6 \pm 0.2‰$, $\delta^{15}N = 11.4 \pm 0.2‰$) provided by SI Science Co. (Saitama, Japan), whose values were determined by the NBS 19 and the International Atomic Energy Agency (IAEA) Sucrose ANU (calibrated against Vienna Pee Dee Belemnite) and IAEA N1 and IAEA N2 (calibrated against AIR) international standards, respectively. The measurement error was estimated as ~0.1‰ for both $\delta^{13}C$ and $\delta^{15}N$ values.

**References**


Bronk Ramsey, C., 1995. Radiocarbon calibration and analysis of stratigraphy: the OxCal program. Radiocarbon. 37, 425–430.

Chen, T., Yu, W.H., Izard, J., Baranova, O. V., Lakshmanan, A., Dewhirst, F.E., 2010. The Human Oral Microbiome Database: a web accessible resource for investigating oral microbe taxonomic and genomic information. Database. 2010, baq013.

Fagernäs, Z., Salazar-García, D.C., Haber Uriarte, M., Avilés Fernández, A., Henry, A.G., Lomba Maurandi, J., Ozga, A.T., Velsko, I.M., Warinner, C., 2022. Understanding the microbial biogeography of ancient human dentitions to guide study design and interpretation. FEMS Microbes. 3, 1–13.

Heaton, T.J., Köhler, P., Butzin, M., Bard, E., Reimer, R.W., Austin, W.E.N., Bronk Ramsey, C., Grootes, P.M., Hughen, K.A., Kromer, B., Reimer, P.J., Adkins, J., Burke, A., Cook, M.S., Olsen, J., Skinner, L.C., 2020. Marine20 - The Marine Radiocarbon Age Calibration Curve (0-55,000 cal BP). Radiocarbon. 62, 779–820.

Hendy, J., Welker, F., Demarchi, B., Speller, C., Warinner, C., Collins, M.J., 2018. A guide to ancient protein studies. Nature Ecology and Evolution. 2, 791–799.

Hughes, C.S., Moggridge, S., Müller, T., Sorensen, P.H., Morin, G.B., Krijgsveld, J., 2019. Single-pot, solid-phase-enhanced sample preparation for proteomics experiments. Nature Protocols. 14, 68–85.

Mackie, M., Rüther, P., Samodova, D., Di Gianvincenzo, F., Granzotto, C., Lyon, D., Peggie, D.A., Howard, H., Harrison, L., Jensen, L.J., Olsen, J. V., Cappellini, E., 2018. Palaeoproteomic profiling of conservation layers on a 14th century Italian wall painting. Angewandte Chemie International Edition. 57, 7369–7374.

Mi, H., Muruganujan, A., Ebert, D., Huang, X., Thomas, P.D., 2019. PANTHER version 14: More genomes, a new PANTHER GO-slim and improvements in enrichment analysis tools. Nucleic Acids Research. 47, D419–D426.

Naito, Y.I., Chikaraishi, Y., Ohkouchi, N., Mukai, H., Shibata, Y., Honch, N. V, Dodo, Y., Ishida, H., Amano, T., Ono, H., Yoneda, M., 2010. Dietary reconstruction of the Okhotsk Culture of Hokkaido,

Japan, based on nitrogen isotopic composition of amino acids: implication for correction of $^{14}C$ marine reservoir effects on human bones. Radiocarbon. 52, 671–681.

Nunoura, T., Chikaraishi, Y., Izaki, R., Suwa, T., Sato, T., Harada, T., Mori, K., Kato, Y., Miyazaki, M., Shimamura, S., Yanagawa, K., Shuto, A., Ohkouchi, N., Fujita, N., Takaki, Y., Atomi, H., Takai, K., 2018. A primordial and reversible TCA cycle in a facultatively chemolithoautotrophic thermophile. Science. 359, 559–563.

Ogura, K., Endo, M., Hase, T., Negami, H., Tsuchiya, K., Nishiuchi, T., Suzuki, T., Ogai, K., Sanada, H., Okamoto, S., Sugama, J., 2023. Potential biomarker proteins for aspiration pneumonia detected by shotgun proteomics using buccal mucosa samples: a cross-sectional case–control study. Clinical Proteomics. 20, 9.

Palmer, K.S., Makarewicz, C.A., Tishkin, A.A., Tur, S.S., Chunag, A., Diimajav, E., Jamsranjav, B., Buckley, M., 2021. Comparing the use of magnetic beads with ultra filtration for ancient dental calculus proteomics. Journal of Proteome Research. 20, 1689–1704.

Reimer, P.J., Austin, W.E.N., Bard, E., Bayliss, A., Blackwell, P.G., Bronk Ramsey, C., Butzin, M., Cheng, H., Edwards, R.L., Friedrich, M., Grootes, P.M., Guilderson, T.P., Hajdas, I., Heaton, T.J., Hogg, A.G., Hughen, K.A., Kromer, B., Manning, S.W., Muscheler, R., Palmer, J.G., Pearson, C., Van Der Plicht, J., Reimer, R.W., Richards, D.A., Scott, E.M., Southon, J.R., Turney, C.S.M., Wacker, L., Adolphi, F., Büntgen, U., Capano, M., Fahrni, S.M., Fogtmann-Schulz, A., Friedrich, R., Köhler, P., Kudsk, S., Miyake, F., Olsen, J., Reinig, F., Sakamoto, M., Sookdeo, A., Talamo, S., 2020. The IntCal20 Northern Hemisphere radiocarbon age calibration curve (0-55 cal kBP). Radiocarbon. 62, 725–757.

Sawafuji, R., Cappellini, E., Fotakis, A.K., Rakownikow, R., Olsen, J. V, Hirata, K., Ueda, S., 2017. Proteomic profiling of archaeological human bone. Royal Society Open Science. 4, 161004.

Sawafuji, R., Saso, A., Suda, W., Hattori, M., Ueda, S., 2020. Ancient DNA analysis of food remains in human dental calculus from the Edo period, Japan. PLoS ONE. 15, e0226654.

Tsutaya, T., Naito, Y.I., Ishida, H., Yoneda, M., 2014. Carbon and nitrogen isotope analyses of human and dog diet in the Okhotsk culture: Perspectives from the Moyoro site, Japan. Anthropological Science. 122, 89–99.

Tsutaya, T., Gakuhari, T., Asahara, A., Yoneda, M., 2017. Isotopic comparison of gelatin extracted from bone powder with that from bone chunk and development of a framework for comparison of different extraction methods. Journal of Archaeological Science: Reports. 11, 99–105.

Tyanova, S., Temu, T., Cox, J., 2016. The MaxQuant computational platform for mass spectrometry-based shotgun proteomics. Nature Protocols. 11, 2301–2319.

Warinner, C., Rodrigues, J.F.M., Vyas, R., Trachsel, C., Shved, N., Grossmann, J., Radini, A., Hancock, Y., Tito, R.Y., Fiddyment, S., Speller, C., Hendy, J., Charlton, S., Luder, H.U., Salazar-García, D.C., Eppler, E., Seiler, R., Hansen, L.H., Castruita, J.A.S., Barkow-Oesterreicher, S., Teoh, K.Y., Kelstrup, C.D., Olsen, J. V, Nanni, P., Kawai, T., Willerslev, E., von Mering, C., Lewis, C.M., Collins, M.J., Gilbert, M.T.P., Rühli, F., Cappellini, E., 2014. Pathogens and host immunity in the ancient human oral cavity. Nature Genetics. 46, 336–344.

Wilkin, S., Ventresca Miller, A., Fernandes, R., Spengler, R., Taylor, W.T.T., Brown, D.R., Reich, D., Kennett, D.J., Culleton, B.J., Kunz, L., Fortes, C., Kitova, A., Kuznetsov, P., Epimakhov, A., Zaibert, V.F., Outram, A.K., Kitov, E., Khokhlov, A., Anthony, D., Boivin, N., 2021. Dairying enabled Early Bronze Age Yamnaya steppe expansions. Nature. 598, 629–633.

Yoneda, M., Uno, H., Shibata, Y., Suzuki, R., Kumamoto, Y., Yoshida, K., Sasaki, T., Suzuki, A., Kawahata, H., 2007. Radiocarbon marine reservoir ages in the western Pacific estimated by pre-bomb molluscan shells. Nuclear Instruments and Methods in Physics Research Section B: Beam Interactions with Materials and Atoms. 259, 432–4

**Supplementary Figures**

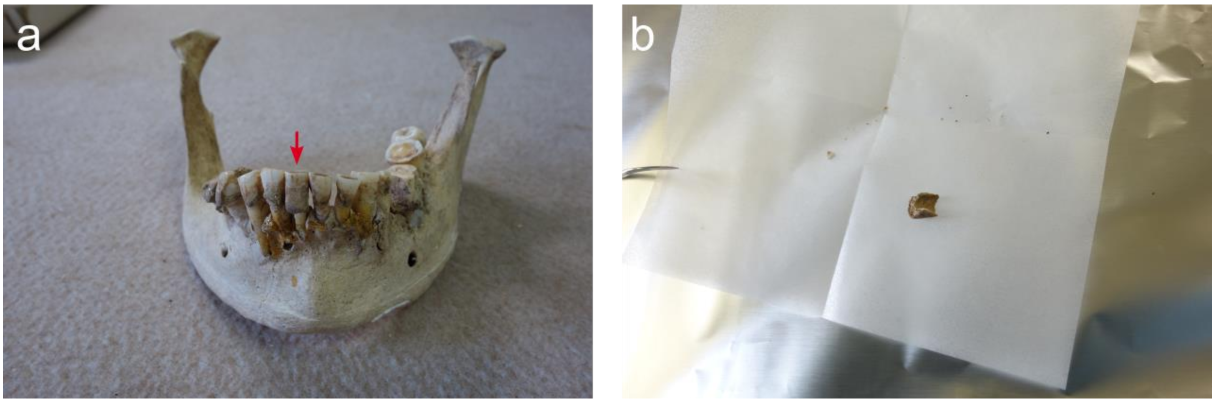


Supplementary Figure S1. Sampled dental calculus of HM2-HA-3. a) Red arrow showing the sampled calculus in situ. b) A part of calculus sampled.

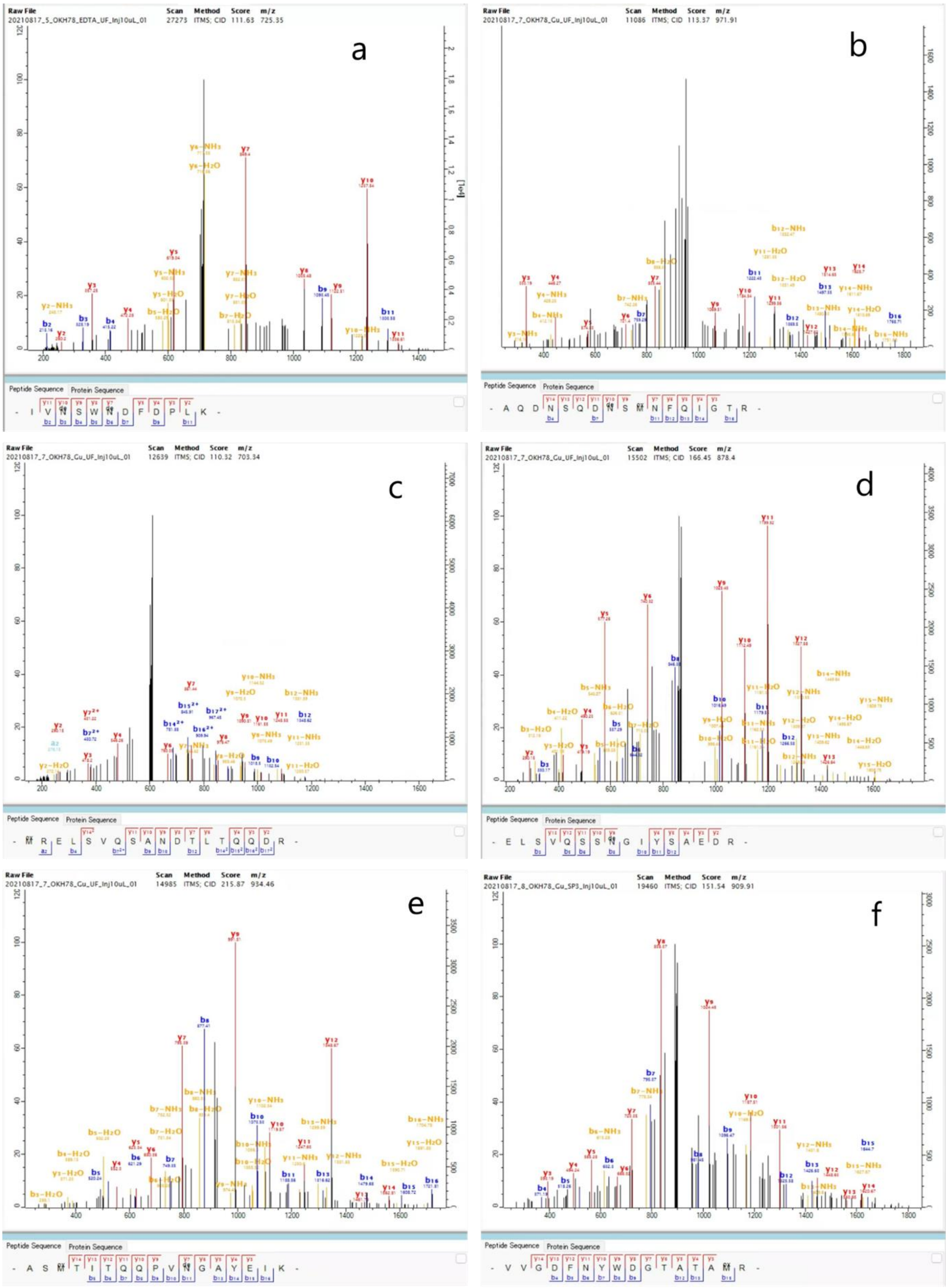


Supplementary Figure S2. MS2 spectra of peptides assigned to periodontitis-related bacterial proteins. a) Inosamine-phosphate amidinotransferase 1, *Actinomyces israelii*, b) Flagellar filament 33 kDa core protein, *Selenomonas sputigena*, c) Flagellin, *Fretibacterium fastidiosum*, d) Flagellar filament 33 kDa core protein, *Treponema denticola*, f) Minor fimbrium subunit Mfa1, *Porphyromonas gingivalis*.

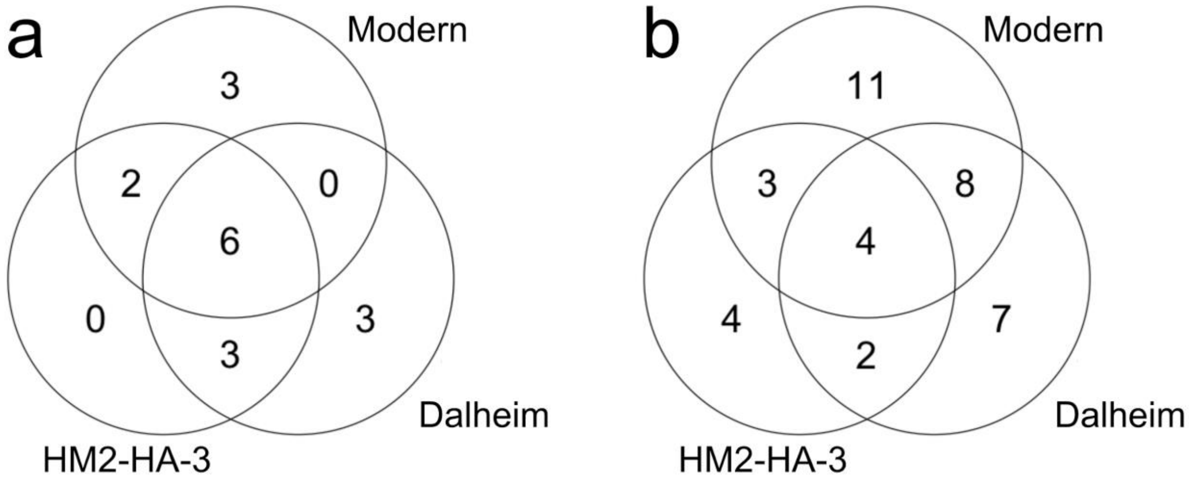


Supplementary Figure S3. Venn diagrams of proteins in a) “defense/immunity” protein class and b) “immune system” biological process identified with PANTHER classification in the ancient dental calculus of HM2-HA-3 (this study), medieval Dalheim, and modern patients (Warinner et al. 2014).

**Supplementary Tables**

Supplementary Table S1. Deamidation rates of asparagine and glutamine in the four fractions of dental calculus of HM2-HA-3. UF: ultrafiltration, SP3: single-pot solid-phase-enhanced sample preparation, S: supernatant, and P: pellet.

Supplementary Table S2. Matched peptides for bio-invasive proteins of periodontitis-related bacterial species. The genera that were matched with BLAST searches of the sequences are shown in the “BLAST” column.

Supplementary Table S3. Human protein groups identified in the ribs of HM2-HA-3.

Supplementary Table S4. Human protein groups identified in the calculus of medieval Dalheim (Warinner et al., 2014).

Supplementary Table S5. Bacterial protein groups identified in the calculus of medieval Dalheim (Warinner et al., 2014).

Supplementary Table S6. Human protein groups identified in the calculus of modern patients (Warinner et al., 2014).

Supplementary Table S7. Bacterial protein groups identified in the calculus of modern patients (Warinner et al., 2014).
